# A Conserved Inhibitory Interdomain Interaction Regulates DNA-binding Activities of Hybrid Two-component Systems in *Bacteroides*

**DOI:** 10.1101/2024.04.25.591157

**Authors:** Rong Gao, Ti Wu, Ann M. Stock

## Abstract

Hybrid two-component systems (HTCSs) comprise a major class of transcription regulators of polysaccharide utilization genes in *Bacteroides*. Distinct from classical two-component systems in which signal transduction is carried out by intermolecular phosphotransfer between a histidine kinase (HK) and a cognate response regulator (RR), HTCSs contain the membrane sensor HK and the RR transcriptional regulator within a single polypeptide chain. Tethering the DNA-binding domain (DBD) of the RR with the dimeric HK domain in an HTCS could potentially promote dimerization of the DBDs and would thus require a mechanism to suppress DNA-binding activity in the absence of stimulus. Analysis of phosphorylation and DNA-binding activities of several HTCSs from *Bacteroides thetaiotaomicron* revealed a DBD-suppression mechanism in which an inhibitory interaction between the DBD and the phosphoryl group-accepting receiver domain (REC) decreases autophosphorylation rates of HTCS-RECs as well as represses DNA-binding activities in the absence of phosphorylation. Sequence analyses and structure predictions identified a highly conserved sequence motif correlated with a conserved inhibitory domain arrangement of REC and DBD. Presence of the motif, as in most HTCSs, or its absence, in a small subset of HTCSs, is likely predictive of two distinct regulatory mechanisms evolved for different glycans. Substitutions within the conserved motif relieve the inhibitory interaction and result in elevated DNA-binding activities in the absence of phosphorylation. Our data suggest a fundamental regulatory mechanism shared by most HTCSs to suppress DBD activities using a conserved inhibitory interdomain arrangement to overcome the challenge of the fused HK and RR components.

**Importance:** Different dietary and host-derived complex carbohydrates shape the gut microbial community and impact human health. In *Bacteroides*, the prevalent gut bacteria genus, utilization of these diverse carbohydrates relies on different gene clusters that are under sophisticated control by various signaling systems, including the hybrid two-component systems (HTCSs). We have uncovered a highly conserved regulatory mechanism in which the output DNA-binding activity of HTCSs is suppressed by interdomain interactions in the absence of stimulating phosphorylation. A consensus amino acid motif is found to correlate with the inhibitory interaction surface while deviations from the consensus can lead to constitutive activation. Understanding of such conserved HTCS features will be important to make regulatory predictions for individual systems as well as to engineer novel systems with substitutions in the consensus to explore the glycan regulation landscape in *Bacteroides*.

## Introduction

Gut microbiota has significant impacts on human health. Perturbation of the complex microbial community has been associated with various diseases, including obesity and diabetes (1–3). Fundamental knowledge of how diets shape the microbiota and how gut microbes utilize nutrients are essential for understanding the mutualistic relationship. Dietary fiber is the main nutrient source for gut microbes. These otherwise undigestible polysaccharides are metabolized to short-chain fatty acids that promote gut health and provide systemic benefits to human hosts (4–6). *Bacteroides*, one of the major bacteria phyla in the distal gut, can utilize a wide variety of polysaccharides. This capability relies on expression of polysaccharide utilization loci (PULs), gene clusters that encode the entire protein machinery for recognition, uptake, hydrolysis, transport and metabolization of complex carbohydrates (7, 8). *Bacteroides* species typically contain a large number of PULs specific for different glycans and thus have developed tight regulatory systems to prevent costly expression of PULs in the absence of their substrates. Hybrid two-component systems (HTCSs) represent one of the major regulatory systems for PULs in *Bacteroides* (8, 9), yet the regulatory mechanisms remain less studied.

HTCSs belong to the family of two-component systems (TCSs), a prevalent prokaryotic signaling scheme involving a phospho-transfer between two conserved protein modules, a histidine kinase (HK) and a response regulator (RR) (10, 11). The HK functions as a dimer and contains multiple enzyme activities that mediate phosphorylation of its cognate RRs via specific HK-RR interactions. Most RRs are transcription regulators containing a DNA-binding domain (DBD). DNA-binding activities are typically regulated by the conserved receiver domain (REC) via phosphorylation-mediated dimerization and/or relief of inhibition (11–13).

HTCS is an unusual class of TCSs with the transmembrane HK fused with the RR transcription regulator in a single polypeptide chain (Fig. 1A). It has been shown that tethering the HK and RR together allows the HK-RR interaction specificity to be promiscuous (14–16) yet individual HTCSs remain insulated from each other because the signaling fidelity can still be maintained due to high local concentrations of cognate partners. This pathway insulation mechanism via tethering does not require co-evolving of interaction specificity as observed in typical TCSs (17, 18) and may enable rapid expansion and evolution of HTCSs for different glycans through domain swapping during gene duplication or lateral transfer events. However, tethering of the HK and RR also presents challenges for regulation because of distinct regulatory mechanisms in HKs and RRs. A phosphorylation-mediated monomer-dimer transition is common for typical RRs to regulate DNA-binding activities. In contrast, an existing dimer is required for typical HKs (10) and transmembrane signaling in HTCSs is through the ligand-induced conformational change within an HK homodimer (19). The dimeric association of HK domains in HTCSs necessarily positions the two tethered DBD domains in close proximity, independent of stimulus, making the monomer-to-dimer RR activation mechanism unlikely to work. DNA-binding activities need to be suppressed within the HTCS dimer in the absence of stimuli. We propose that suppression of DNA-binding activities is likely to occur via interdomain interactions between DBD and other HTCS domains.

**FIG 1.**
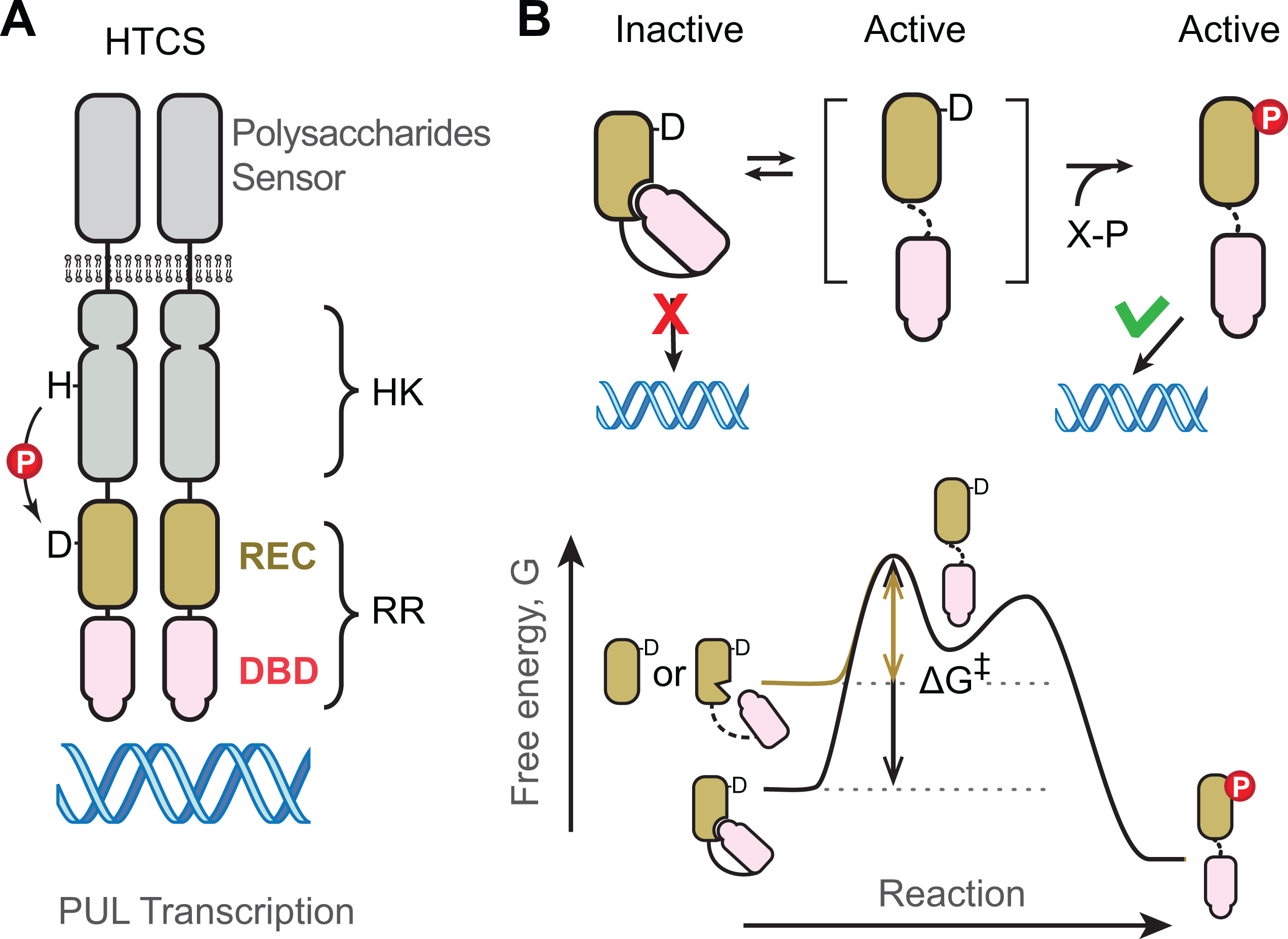
Schematic of HTCS regulation. (A) Domain organization of HTCSs. Activation of HTCSs by polysaccharides is through the functional dimer of histidine kinase domains. The DNA-binding activities need to be regulated within the dimer and further mediated by phosphorylation of the REC domain. (B) Effects of interdomain interaction on phosphorylation kinetics and DNA binding activities. Inhibitory interaction shifts the conformational equilibrium, increases the free energy barrier, and slows down phosphorylation kinetics. Isolated REC domains or RR variants with disrupted interaction (brown line) will have a smaller energy barrier for phosphorylation than RRs with the inhibitory interaction (black line).

Inhibitory interdomain interactions between the DBD and REC domains have been described for many RRs (12, 13, 20). In addition to interactions that bury surfaces of the REC domains that mediate dimerization, the DNA-recognition helix of DBDs can be buried at the REC-DBD interface or positioned unfavorably for binding to DNA. Owing to the plasticity of REC domains, interaction surfaces and domain arrangements can differ greatly for RR subfamilies with different DBDs or even within the same subfamily of RRs (11, 13, 20, 21). Nevertheless, regulation relies on the phosphorylation-activated switch of allosteric conformations of the REC domain. The REC domain exists in equilibrium between inactive and active conformations, with phosphorylation occurring only within the active conformation (22–24) and shifting the equilibrium to the active state to allow DNA-binding (Fig. 1B). Inhibitory interactions can stabilize the inactive conformation and prevent RR activation in the absence of phosphorylation. On the other hand, this stabilization can impact phosphorylation kinetics by lowering the free energy of the inactive conformation and increasing the free energy barrier for the phosphorylation reaction (Fig. 1B). Phosphorylation experiments using small-molecule phospho-donors, such as acetyl-phosphate and phosphoramidate (PAM), have shown a slower phosphorylation speed for intact RRs with linked REC and DBD domains than for the isolated REC domain alone (13). Thus, comparing the phosphorylation kinetics of the REC domain and the RR fragment (REC+DBD) of HTCSs can potentially infer whether an inhibitory interaction exists between the REC and DBD domains.

Here we investigate phosphorylation kinetics of RR fragments of several HTCS proteins from *Bacteroides thetaiotaomicron* (*B. theta*). All displayed slower phosphorylation kinetics with the DBD linked than with the REC domain alone, suggesting an inhibitory interaction between the REC and DBD domains. Sequence conservation analyses and structure predictions indicate a highly conserved motif at the α3-β4 loop of the REC domain potentially involved in the REC-DBD interactions. Substitutions within this motif disrupt the inhibitory interaction and allow the DBD to bind DNA in the absence of RR phosphorylation. Collectively, our findings uncover a highly conserved mechanism for regulating DNA-binding activities of HTCSs in *Bacteroides*.

## Results

### Phosphorylation kinetics of RRs and RECs

The defining feature of a REC domain is its highly conserved phosphorylation site that makes it catalytically competent for autophosphorylation once bound to a phospho-donor, such as phosphoramidate (PAM). Although the contribution of phosphorylation by small molecules is not always physiologically significant, the phosphorylation kinetics reflect the activation energy for catalysis and may reveal potential interdomain interactions. An inhibitory interaction between the REC and DBD domain can slow down the phosphorylation rate. A few HTCS proteins from *B. theta* with known glycan substrates, including BT4124 (25), BT4663 (19), BT3334 (26) and BT1754 (27), were selected for in vitro phosphorylation analyses.

Phosphorylation of purified RR fragments (REC+DBD) and REC domains upon addition of PAM was analyzed using Phos-tag gels (28) that separate phosphorylated and unphosphorylated proteins (Fig. 2A, Fig. S1 and S2). All RECs were observed to be phosphorylated faster than corresponding RRs. For example, significant phosphorylation was observed for BT4124-REC at 0.2 min after addition of 20 mM PAM while it took longer, 0.7-2 min, for BT4124-RR to reach a similar level of phosphorylation (Fig. 2A). Fractions of phosphorylated proteins can be quantified by measuring the band intensities of phosphorylated proteins relative to the total intensities of both unphosphorylated and phosphorylated proteins, yielding the kinetic curves shown in Fig. 2B. Phosphorylation reached a steady level at later time points, indicating a balance of autophosphorylation and autodephosphorylation rates. The steady-state level of REC phosphorylation did not reach full phosphorylation, but was higher than that of the RR, consistent with a higher autophosphorylation rate. More importantly, the half time of phosphorylation for BT4124-REC is 0.3 min, much shorter than the half-time of 1.5 min for BT4124-RR (Fig. 2B). A similar trend with shorter half-times for RECs than for RRs was also observed for other HTCSs (Fig. S2A).

**FIG 2.**
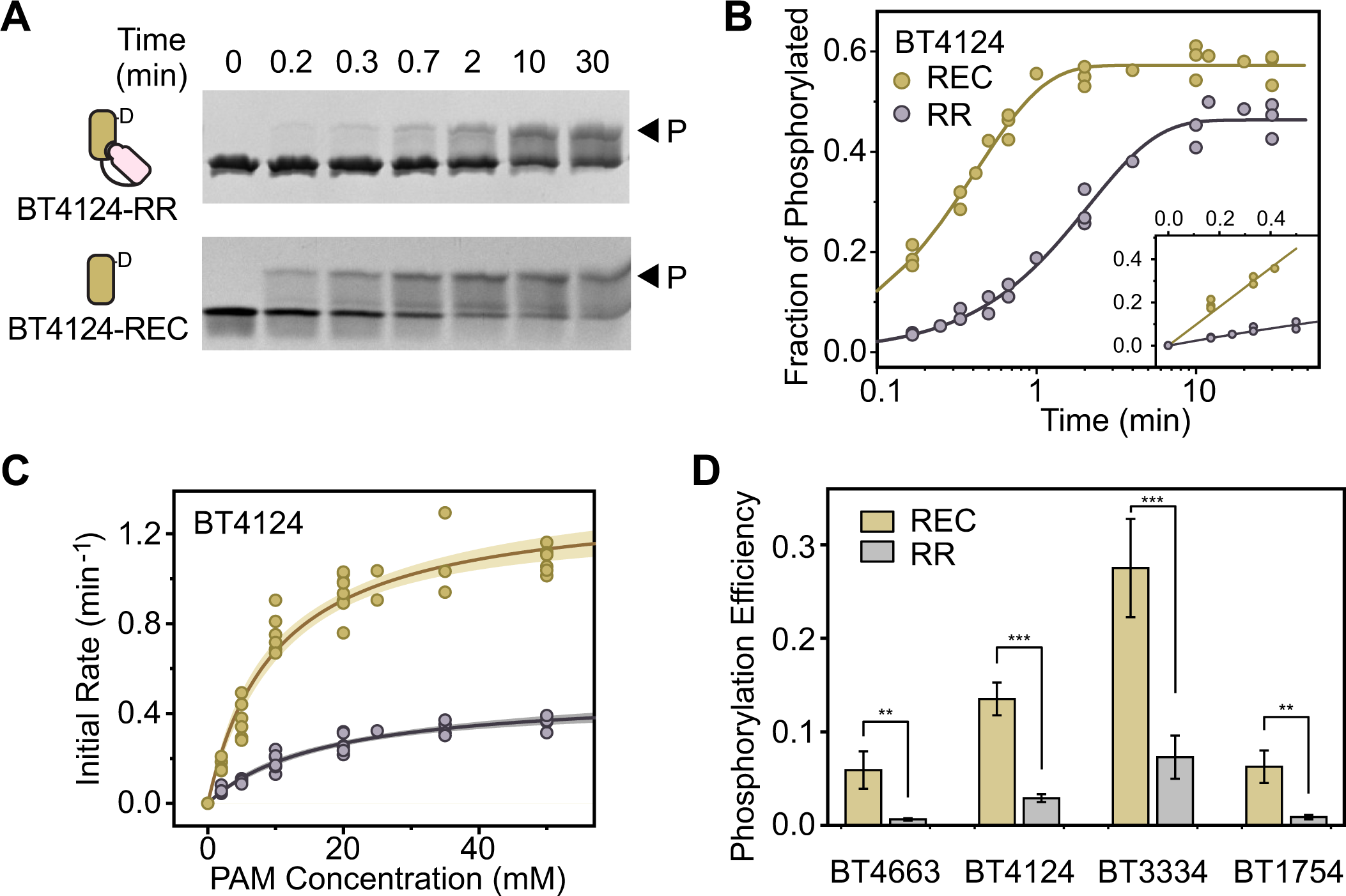
Isolated REC domains autophosphorylate faster than HTCS RRs with linked DNA binding domains. (A-C) Autophosphorylation kinetics of BT4124. Phosphorylation of BT4124 proteins were analyzed using Phos-tag gels at indicated times after addition of 20 mM PAM (A). One representative example is shown for each protein. Fractions of phosphorylated proteins from (A) were quantified to derive the exponential trendline of phosphorylation in (B). Initial rates of phosphorylation were calculated from early stages of reaction (inset in B). (C) Kinetic characterization of BT4124 phosphorylation. Independent measurements of initial rates are plotted as circles. Solid lines indicate individual fits with the Michaelis-Menten equation and the shaded areas represent the 95% confidence intervals. (D) Comparison of phosphorylation efficiency between RRs and REC domains across different HTCS proteins. Error bars indicate the standard deviation calculated as described in Methods.

For more rigorous comparison of phosphorylation kinetics, initial rates were derived from early stages of phosphorylation (Fig. 2B inset). Across all PAM concentrations, phosphorylation rates of HTCS-RR proteins appeared to be lower than those of corresponding RECs (Fig. 2C, Fig. S1 and Fig. S2). Initial rates were fitted with a hyperbola function based on the Michaelis-Menten kinetics to derive kinetic parameters (Table S1). RECs have higher *k_cat_* and lower *K_m_* values than RRs, suggesting that the linked DBD domain may impact the phosphorylation catalysis, and potentially the binding of PAM. Rather than scrutinizing individual parameter values and model details that depend greatly on the complexity of the phosphorylation reaction pathway, we chose to focus on the kinetic parameter, *k_cat_*/*K_m_*, termed the phosphorylation efficiency (Supplementary Text). The *k_cat_*/*K_m_*value is inversely related to the free energy barrier of transition states in various simple or complex enzyme pathways (29). The *k_cat_*/*K_m_* values of all four tested HTCS-RRs are significantly lower than those of corresponding isolated RECs (Fig. 2D). Phosphorylation efficiencies are suppressed by the presence of the DBD domain, ranging from ∼1/10 for BT4663-RR to ∼1/4 for BT3334-RR (Table S1). Lower efficiencies for RRs than for RECs suggest that the linked DBD domain in individual HTCS-RR fragments increases the activation energy for the phosphorylation reaction, which can result from inhibitory interactions between the REC and DBD domains.

### Predicted domain interaction of REC and DBD

To explore the potential inhibitory interaction, AlphaFold 2 was used to predict the structural arrangements of the REC and DBD domains (30, 31). BT4124-RR is used as an example to illustrate the confidence of prediction. The five top-ranked predicted structures have identical domain arrangements (Fig. S3A) and the entire structure except for the loop connecting the REC and DBD, the N- and C-termini has a high confidence score (Fig. S3B). Fig. 3A shows the 1^st^- ranked predicted structure of BT4124-RR. The REC domain has the typical (βα)_5_ fold with a long α5 helix extending toward the DBD. The DBD belongs to the AraC/XylS family of helix-turn-helix (HTH) domains, also known as HTH18 in PFAM (PFAM id, PF12833), and contains two DNA-recognition helices (*gold*) within two HTH motifs.

**FIG 3.**
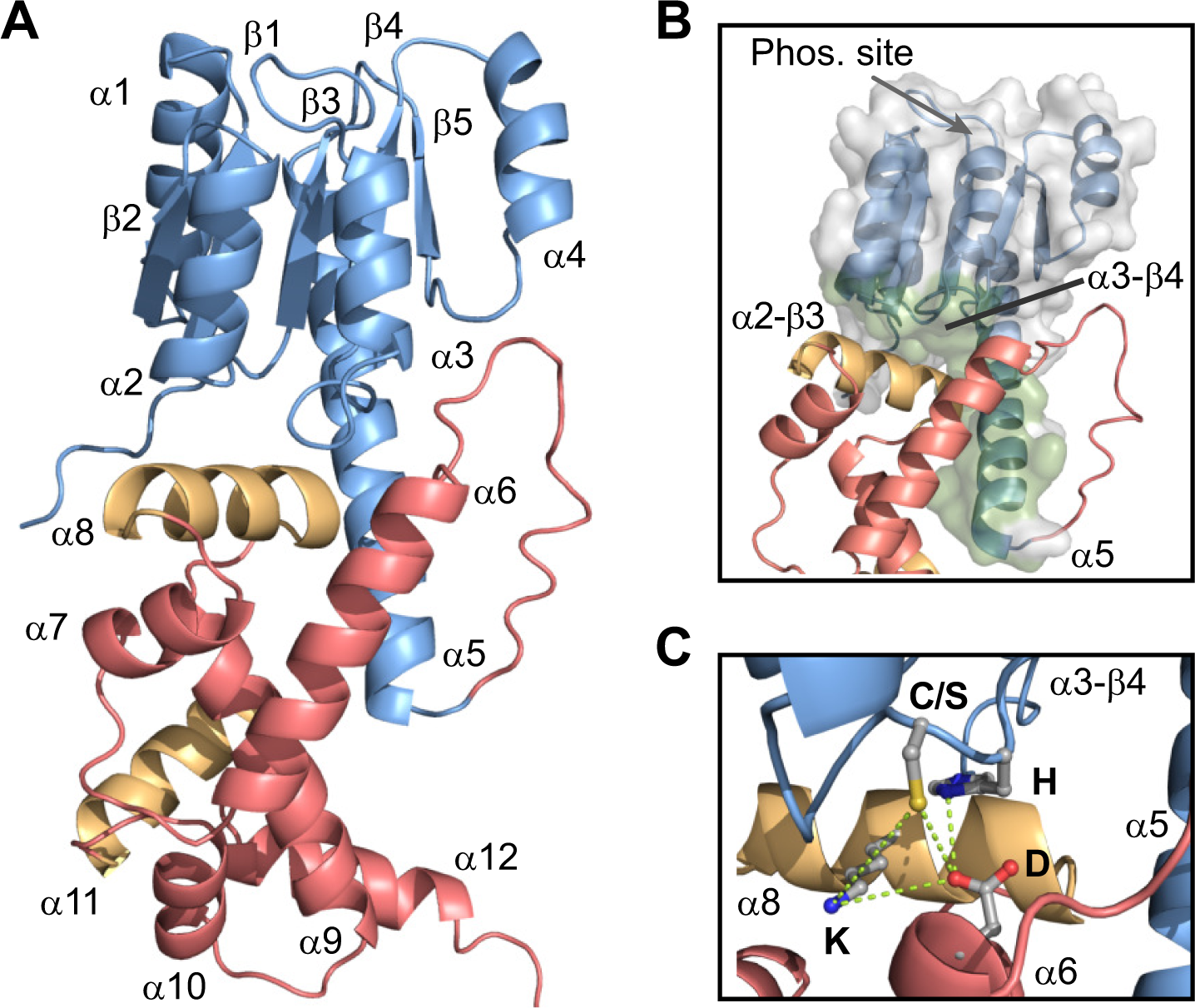
Predicted structure of an HTCS RR suggests interaction between the REC and DBD domains. (A) Ribbon view of the BT4124-RR structure predicted by AlphaFold 2. The highest ranked structure is shown. The REC domain is colored blue and the DBD domain is colored red with the two DNA-recognizing helices highlighted in gold. (B) The interaction surface (green) of the REC domain that contacts the DBD. (C) Predicted contact residues at the REC-DBD interface.

The DBD is predicted to make extensive contacts with the REC domain. The interaction surface involves the extended α5 helix and various α-β loops (Fig. 3B), including α2-β3 and α3- β4, which are at the opposite face of the domain from the phosphorylation site located within the β-α loops. One of the two DNA-recognition helices, α8, is buried at the interface, directly packed against the α-β loops of the REC domain. Superimposing the predicted structure of BT4124- DBD with the structure of DNA-bound HTH18 family member MarA (32) indicates a significant clash of DNA with the REC domain (Fig. S3C). Thus, this domain arrangement of REC and DBD is likely to preclude DNA binding and the interdomain interaction may play an inhibitory role for RR activation. The observed interface may also stabilize a specific conformation of the REC domain and contribute to the observed suppression of phosphorylation rates by DBDs.

### Conserved interaction residues at the predicted interface

It has been shown that REC-DBD interaction surfaces in typical RRs often involve hydrogen bonds between polar or charged residues at the center of interaction surface (13). For example, interdomain hydrogen bonds between Tyr and Asp, Tyr and Asn, or Asp and Gln residues have been described for DrrB, PrrA and VraR, respectively (12, 21, 33). A few similar residues at the heart of the predicted interaction surface of BT4124, including C^1262^ and H^1263^ in the α3-β4 loop of REC, D^1344^ in α6 of the DBD, and K^1383^ in the recognition helix α8, are in close proximity with each other (Fig. 3C). H^1263^ is within hydrogen bond distance of D^1344^ while C^1262^ is close to both D^1344^ and K^1383^, all of which may provide a hydrogen-bonding network to anchor the interaction. To investigate whether these residues are conserved in all HTCSs, sequence conservation was evaluated for 6908 HTCS proteins from *Bacteroides* (Fig. 4 and Fig. S4). Total information contents (ICs) at each position were normalized to the maximum of IC and used to represent the degree of sequence conservation. Not surprisingly, residues constituting the phosphorylation/phosphotransfer active site (asterisks in Fig. 4) at multiple β-α loops are among the most conserved regions in the REC family as well as the HTCS-REC subgroup.

**FIG 4.**
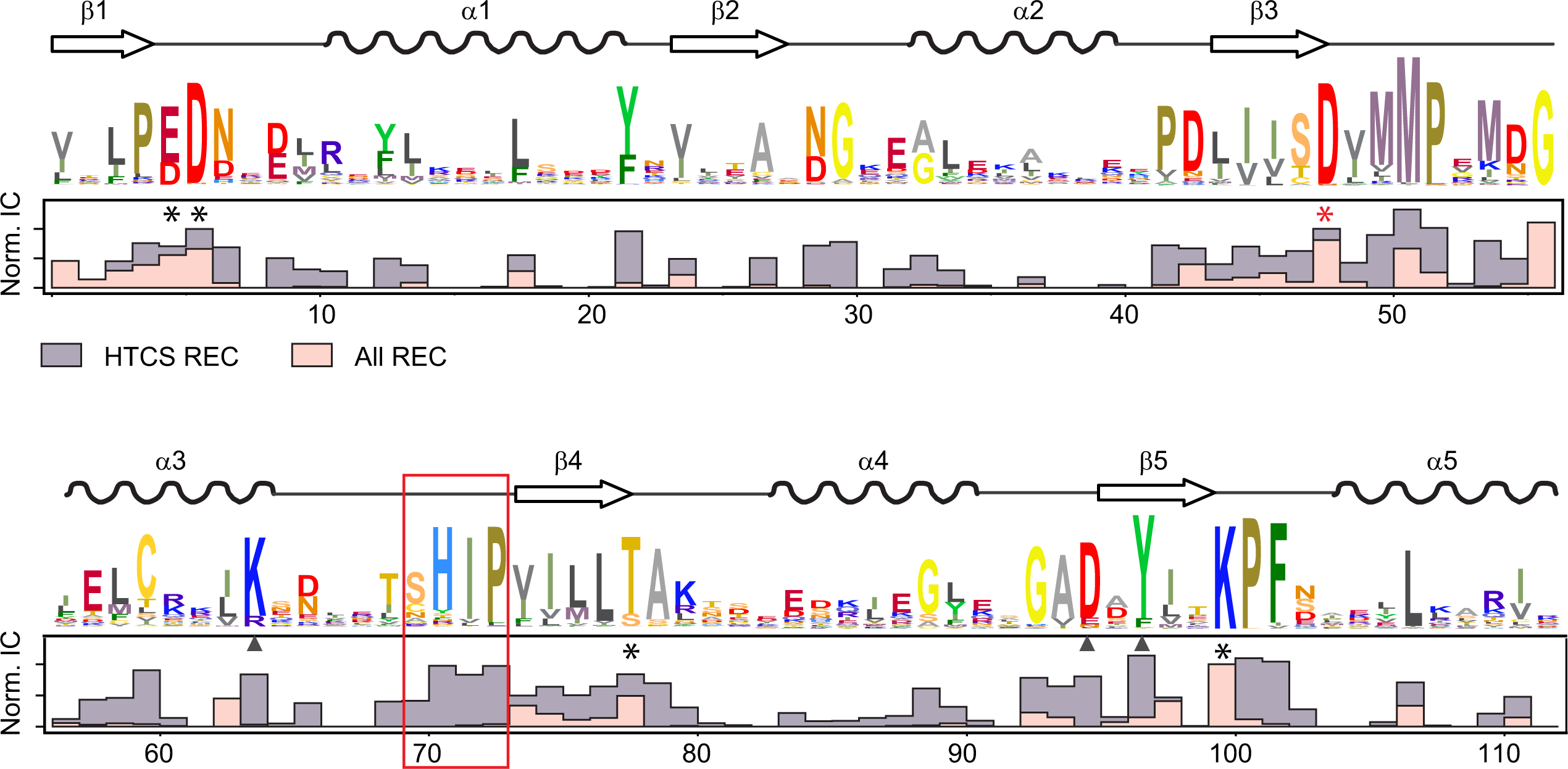
HTCS-REC domains have highly conserved regions not observed in typical REC domains. Sequence conservation of HTCS-REC domains is illustrated by sequence logos obtained from the profile hidden Markov model of 6908 HTCS proteins in *Bacteroides*. Secondary structural elements are shown above the logos. Bar graphs below compare the normalized information contents (ICs) between HTCS-RECs (gray) and the entire REC family from PFAM PF00072 (pink). In addition to highly conserved phosphorylation site residues (asterisks) at the β-α loops, HTCS-RECs display sequence conservation in other regions, such as the α3-β4 loop (red box) and localized positions at α3 and β5 (triangles).

In contrast to the whole REC family, which has little sequence conservation in regions other than the phosphorylation site, HTCS-RECs display exceptionally high sequence conservation in several regions not associated with phosphorylation. These conserved regions include the β5 strand and its adjacent loops, which share some sequence similarity with the OmpR/PhoB subfamily of RRs (34), the α3 helix, and the α3-β4 loop. A sequence motif (S/CHIP) in the α3-β4 loop is among the most conserved stretches of sequence in HTCS-RECs (red box in Fig. 4). At the first position of this conserved motif, 84% of HTCS sequences have a Ser or Cys (S/C) and more than 90% of sequences have the His, Ile and Pro residues (HIP) at the latter three positions. The first two residues (S/CH) correspond to the potential hydrogen-bond-forming residues observed at the predicted REC-DBD interface while the latter two residues (IP) may function to ensure proper loop orientation for contacting the DBD. Correspondingly, the D residue in helix α6 and K residue in helix α8 at the contact surface are also highly conserved only in HTCS-DBDs and not in other HTH18 family members (Fig. S4). Given the sequence conservation pattern for these residues (S/CHIP-DK), the predicted domain arrangement and inhibitory interaction involving these contact residues may represent the basis for a conserved mechanism for regulating the DNA-binding activities.

### Correlation between the conserved motif with the REC-DBD domain orientation

Structural prediction of the RR fragment was performed for all 32 HTCSs in *B. theta* to investigate whether HTCSs may share a conserved domain orientation for interdomain inhibition. Most HTCS-RR structures have a similar domain arrangement as BT4124 with the root mean square distance (RMSD) values below 4 Å (Fig. 5A). Among them are BT3334 and BT4663, with the DNA recognition helix α8 packing against the α3-β4 loop at the REC-DBD interface (Fig. S5A). Similar to BT4124, this REC-DBD interface may account for the observed phosphorylation suppression in BT3334 and BT4663. BT1754-RR is predicted to have a distinct domain arrangement (RMSD, 17.7 Å) but the pTM confidence score, reflecting the topological accuracy (30, 35), is not high. Because the lack of the linked HK domain may impact the interaction or spatial access of REC and DBD, the entire cytoplasmic portion of HTCS (HTCS- cyto), containing both HK and RR, is used to predict the dimeric structure of HTCS-cyto. The majority of HTCS-cyto proteins (25/32), including BT1754-cyto (Fig. S5A), are predicted to have a similar domain orientation as BT4124. Strikingly, all these HTCSs with a similar REC- DBD interface also have the fully conserved motif (S/CHIP-DK) while HTCSs with sequences deviating from the consensus have different domain arrangements with large RMSD values (Fig. 5A). It appears that this conserved motif is predictive of the inhibitory interdomain contact that suppresses the DNA binding activity of DBDs.

**FIG 5.**
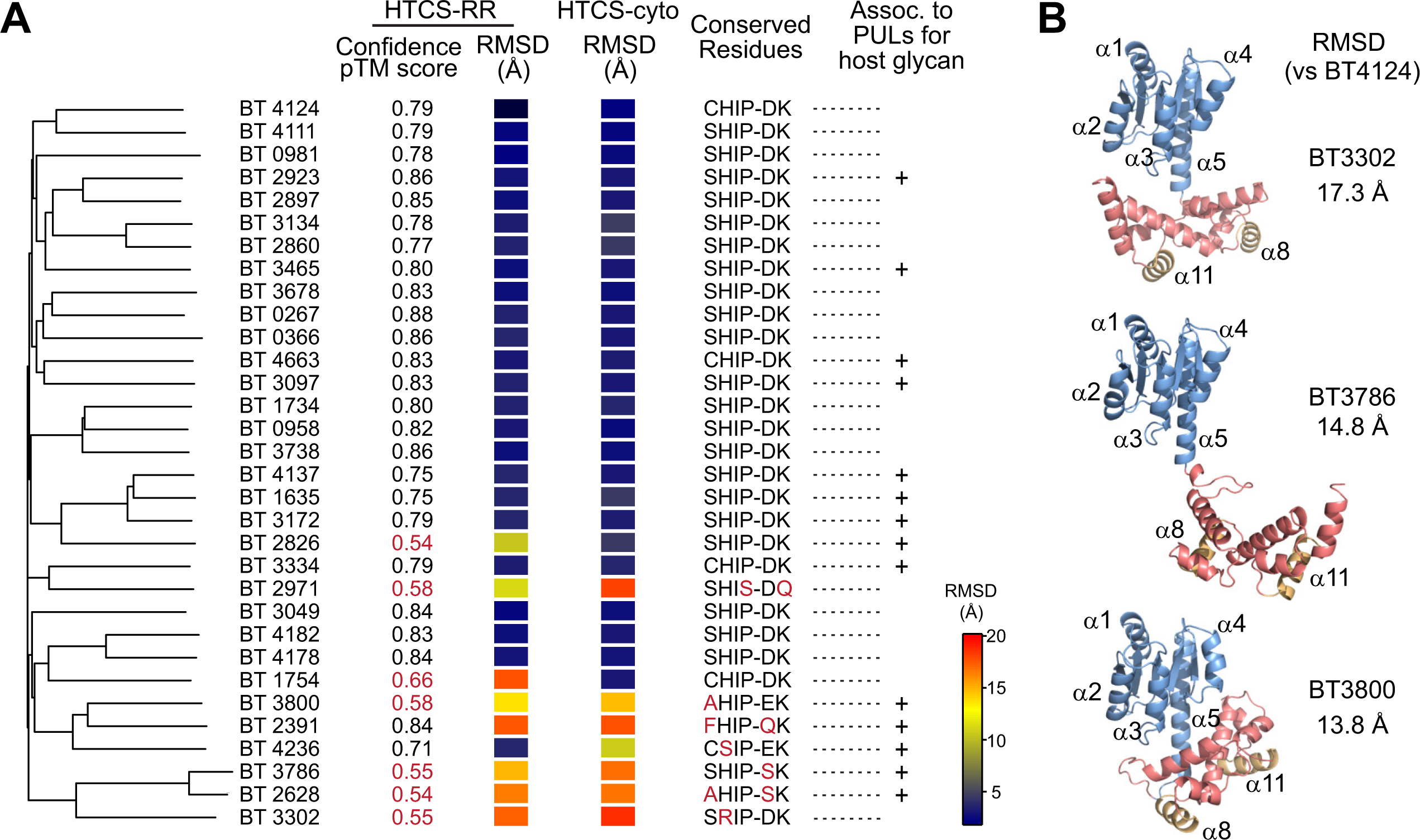
The conserved sequence motif correlates with the domain orientation in HTCSs from *B. theta*. (A) Comparison of predicted structures of HTCS-RR or HTCS-cyto with BT4124-RR. Sequences of the RR fragment of 32 HTCSs from *B. theta* were aligned with Clustal Omega to generate the phylogenetic tree using the neighbor-joining method and default parameters. RMSD values reflect the similarity of domain orientation to that of BT4124-RR. The “+” signs highlight the HTCSs that are within PULs involved in host glycan degradation (36). (B) Diverse domain orientations in HTCSs that have motif sequences that deviate from the consensus. Three representative HTCSs are shown. Predicted structures of other HTCSs, such as BT2391, BT2628, BT2971 and BT4236, are shown in Fig. S5 and S6.

For HTCSs with deviations in the consensus motif, a wide variety of domain positions have been observed for predicted structures. The relative positioning of DBD to REC can be different for different HTCS proteins (Fig. 5B), different fragments of the same protein, HTCS-cyto and HTCS-RR (Fig. S5B) or even different ranked predictions of the same HTCS fragment (Fig. S6). Many of the structures, such as BT2391, BT3302 and BT3786 (Fig. 5B and S5B), have small or nearly no contact between the REC and DBD domains. One extreme example is BT2628 that shows five different orientations with little interdomain contact for five ranked predictions with comparable confidence scores (Fig. S6). Interdomain accuracy of the AlphaFold prediction correlates with the pTM score and will decrease if domains are mobile relative to each other (30). Most HTCS-RRs that show non-conserved domain positioning also have lower pTM scores (highlighted red in Fig. 5A) than those with the conserved orientation. Plasticity of the domain orientation may be a sign of no restrictive interaction between REC and DBD. Sequence deviations from the consensus can result in loss of hydrogen bonds at the REC-DBD interface, which relieves the inhibitory interaction. Moreover, these non-consensus HTCSs appear to be phylogenetically close (Fig. 5A) and many are associated with PULs that degrade host glycan (36). An alternative regulatory strategy distinct from the conserved inhibition mechanism may have evolved for this small group of HTCSs.

### Effects of interface substitutions on phosphorylation

Structural prediction and sequence conservation suggest the importance of the S/CHIP-DK motif. If they contribute substantially to the stability of the interface, substitution of these residues might disrupt the REC-DBD interaction. Because the D and K residues in the DBD are located near or within the DNA-recognition helix α8, alteration of the two residues may have complex pleiotropic effects on DNA recognition. On the other hand, the S/CH residues at the α3-β4 loop of REC are far from the phosphorylation site and would be expected to have less direct impact on REC phosphorylation and more specific effects on REC-DBD interactions. The C^1262^H^1263^ residues in BT4124-RR were altered to A^1262^G^1263^ or A^1262^D^1263^ to disrupt the potential hydrogen bonds and the substituted proteins were named BT4124-RR^AG^ and BT4124-RR^AD^, respectively. Replacing H^1263^ with a D residue in BT4124-RR^AD^ introduces a negatively charged residue near D^1344^ at α6, which may have a greater impact on disrupting the inhibitory interaction.

Both BT4124-RR^AG^ and BT4124-RR^AD^ displayed faster phosphorylation kinetics (Fig. 6A), suggesting a relief of the inhibitory interaction between the REC and DBD domains. At 20 mM PAM, the phosphorylation half-time is 0.4 min for BT4124-RR^AG^ and 0.3 min for BT4124-RR^AD^, similar to the half-time of 0.3 min for BT4124-REC that is not inhibited by the DBD. Phosphorylation efficiencies of the two substituted proteins are significantly higher than BT4124-RR, with BT4124-RR^AD^ indistinguishable from BT4124-REC (Fig. 6B and Fig. S7). This implies that the AD substitution completely abolishes the interdomain inhibition. BT4124- RR^AG^ appears to have slightly lower phosphorylation efficiency than BT4124-RR^AD^ with a *p*- value at 0.04, suggesting partial relief of inhibition, different from BT4124-RR^AD^. Nevertheless, both AG and AD substitutions of the contact residues greatly accelerate the phosphorylation.

**FIG 6.**
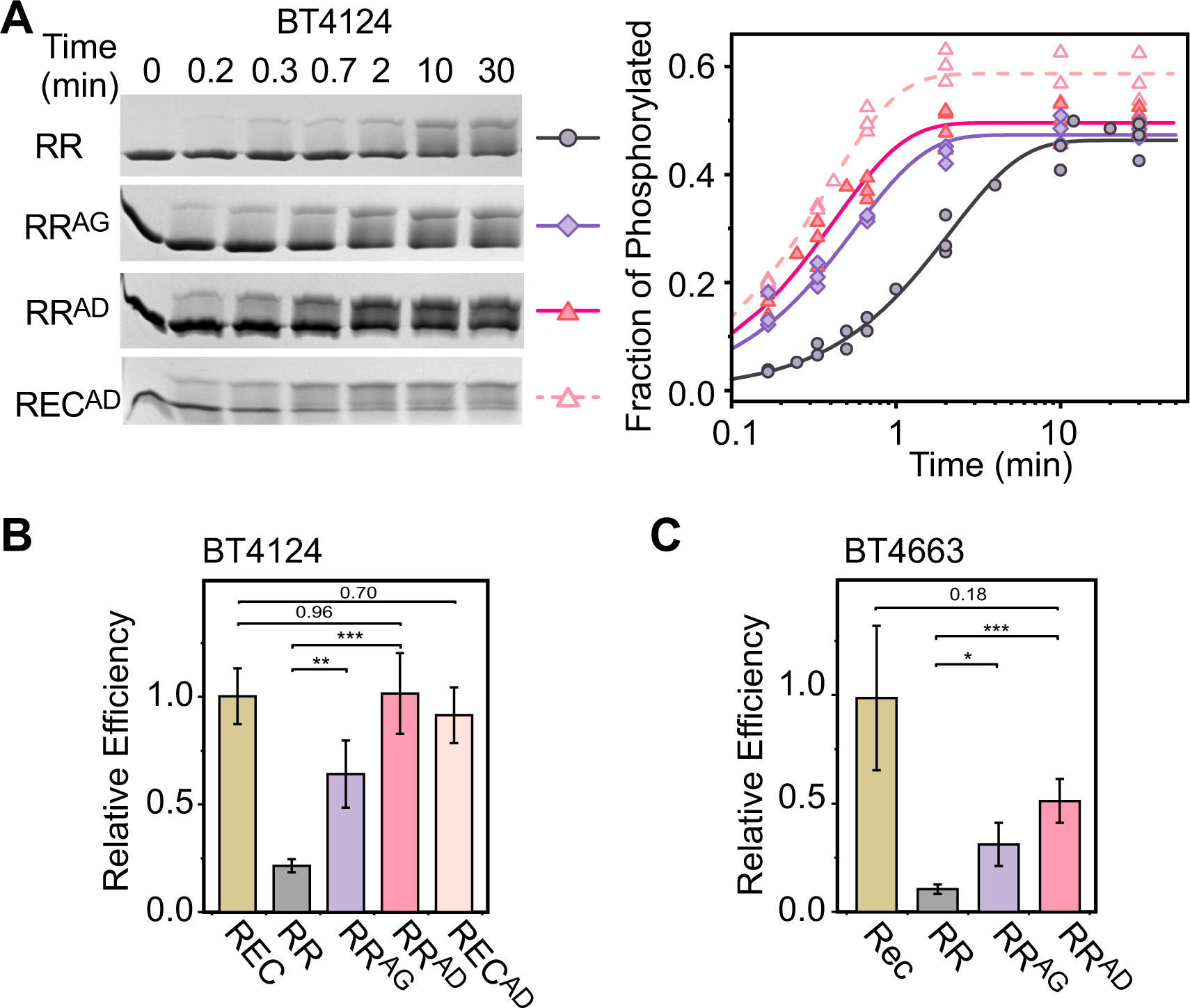
REC-DBD interface substitutions relieve inhibition and accelerate phosphorylation. (A- B) Autophosphorylation kinetics of BT4124 domains and corresponding interface variants. Phos- tag gels (A) were quantified to track the fraction of phosphorylated proteins (B) at indicated times after addition of 20 mM PAM. (C-D) Comparison of phosphorylation efficiency for RR and REC-DBD interface variants of BT4124 (C) and BT4663 (D). Error bars indicate the standard deviation calculated as described.

To exclude the possibility that the substitutions have a direct effect on the phosphorylation site, phosphorylation kinetics were compared for BT4124-REC and BT4124-REC^AD^. The AD substitution in isolated REC domain did not alter the kinetic profile (Fig. S7B) and the phosphorylation efficiency of BT4124-REC^AD^ is not significantly different from that of BT4124-REC. Therefore, substitutions of the contact residues in BT4124-RR are not likely affecting the phosphotransfer active site directly, but are rather relieving the inhibitory interactions, allowing for faster phosphorylation. A similar pattern of phosphorylation efficiencies was also observed in BT4663-RR (Fig. 6C and Fig. S8). Both AG and AD substitutions increased the catalytic efficiency. Taken together, the CH residues appear to be essential for maintaining the inhibitory interaction between the REC and DBD domains.

### Effects of interface substitutions on DNA-binding

Acceleration of PAM phosphorylation kinetics for proteins containing substitutions of conserved interface residues indicates a relief of interdomain inhibition. The direct consequence of disrupting the inhibitory interaction will be the loss of regulation of DBD activity, relieving sequestration of the recognition helix and allowing the DBD to bind DNA in the absence of phosphorylation. Therefore, we examined the DNA-binding activities of HTCS-RRs and corresponding interface substituents using electrophoretic mobility shift assays (EMSAs) (Fig. 7). Promoter DNA fragments from *bt4114* for BT4124-RR and *bt4662* for BT4663-RR were used for DNA-binding studies because *bt4114* and *bt4662* promoters drive transcription of *susCD* homologs of corresponding PULs and represent the regulatory targets of BT4124 and BT4663, respectively (9).

**FIG 7.**
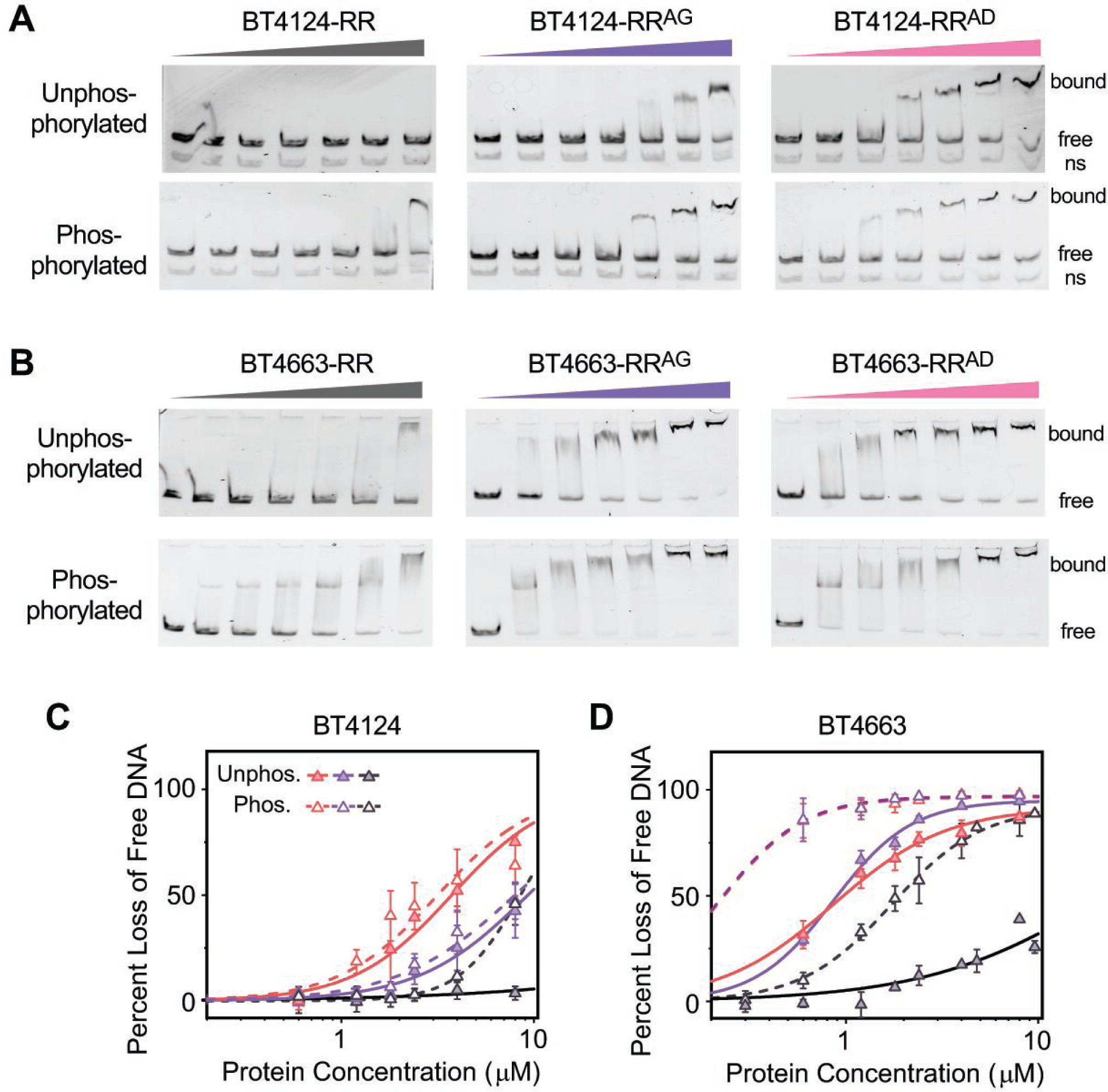
REC-DBD interface substitutions relieve inhibition and allow DNA-binding. (A-B) Binding of HTCS-RR proteins to the DNA detected by EMSAs. EMSAs were done with PCR-generated fluorescent DNA fragments in the presence of 0, 0.6, 1.2, 1.8, 2.4, 4, 8 μM concentrations of the indicated proteins. Promoter DNA from *bt4114* was used for BT4124 proteins (A) and promoter DNA from *bt4662* was used for BT4663 proteins (B). A non-specific DNA fragment (“ns”) present in all lanes in (A) did not show any shift. (C-D) Comparison of DNA-binding curves of HTCS RR proteins. Percent decreases of band intensities for the free unbound DNA were quantified. RR, RR^AG^ and RR^AD^ are colored black, violet and pink, respectively. Data are shown as mean ± SD from at least three independent experiments. Solid (unphosphorylated) and dashed (phosphorylated) lines represent data fitted with the Hill equation.

DNA-binding activities of HTCS-RR proteins depend on phosphorylation. In the absence of phosphorylation, BT4124-RR did not bind to DNA while phosphorylation enabled DNA-binding at high concentrations of BT4124-RR (Fig. 7A, *left*). Phosphorylation also greatly enhanced the DNA-binding activities of BT4663-RR (Fig. 7B, *left*). Quantification of the free unbound DNA was used to generate binding curves (Fig. 7C and 7D). Accurate determination of the DNA affinity was not possible without simultaneous measuring the protein phosphorylation levels. Instead, apparent affinity was derived from the binding curves for qualitative assessment of DNA-binding activities. Phosphorylated BT4663-RR has an apparent *K_D_* of 1.7 μM for binding DNA, much stronger than the unphosphorylated protein, which has an apparent *K_D_* of >22 μM.

Interface variants containing the substitutions AG and AD showed significant DNA affinity in the absence of phosphorylation, supporting the hypothesis that disruption of the REC-DBD interaction relieves the inhibition of DBD activity. Both BT4124-RR^AG^ and BT4124-RR^AD^ bound more DNA in the absence of phosphorylation than BT4124-RR (Fig. 7A and 7C).

Comparing the two variants, BT4124-RR^AG^ binds to DNA less tightly than BT4124-RR^AD^, consistent with the partial relief of inhibition suggested by phosphorylation efficiencies shown in Fig. 6B. Phosphorylation of the BT4124-RR variants did not alter the DNA-binding activities significantly. In contrast, phosphorylation of BT4663-RR^AG^ and BT4663-RR^AD^ greatly enhances the affinity for DNA (Fig. 7D). The differences between BT4124-RR and BT4663-RR may reflect differences in contribution of peripheral residues at the REC-DBD interface or other phosphorylation-induced structural changes. Despite the differences in effects of phosphorylation between BT4663 and BT4124 variants, interface substitutions AG and AD in BT4663-RR also relieve DBD inhibition and allow the DBD to bind DNA in the absence of phosphorylation. The apparent *K_D_* is 0.9 μM for both BT4663-RR^AG^ and BT4663-RR^AD^, much less than that of unphosphorylated BT4663-RR. A common mechanism involving the conserved contact residues at the α3-β4 loop of REC appears to be important to suppress DNA binding activity of the DBD.

## Discussion

In addition to having fitness benefits in glycan utilization (37, 38), PULs also carry considerable fitness costs depending on the glycan nutrient environments (39). Gratuitous expression of PULs can be detrimental to cell survival. Preventing futile expression is likely to be as important as activating PULs in response to dietary and host glycans. Accurate regulation of PULs is essential for prioritized utilization of different polysaccharides by *Bacteroides* species, which impacts the competitive and mutual relationships among gut microbes (40–42). HTCSs regulate transcription of PULs via modulating the DNA-binding activities of the AraC-type DBD. It has been shown that placing two DBDs of AraC adjacent to each other by fused dimerization domains results in transcription activation (43), thus tethering DBDs with the HK dimer in HTCSs may give rise to unnecessary activation of PULs and requires a DBD-suppression strategy. Our studies demonstrate phosphorylation-dependent DNA binding activities and reveal a highly conserved molecular mechanism that inhibits activities of HTCS-DBDs via an interdomain interaction with the REC domain.

### Interaction plasticity between the DBDs and their regulatory domains

Inhibition of DNA-binding activities is a common regulatory strategy utilized by RRs (11) or other transcription regulators, such as those containing a ligand-binding domain and an AraC-type DBD (44, 45). These regulatory proteins usually exist in equilibrium between an active conformation competent for transcription activation and an inactive form whose activities are repressed by interdomain interactions. Either phosphorylation of REC or ligand-binding in the AraC-type regulators shifts the conformation dynamics between the two states. Structures of the inactive form of these regulators show great variations in contact surfaces involved in the inhibitory interaction. For example, three AraC-type regulators with distinct ligand-binding regulatory domains, ToxT (44), CdpR (46) and XylR (47), display three different domain arrangements. In ToxT, the regulatory domain interacts the DBD at the helix connecting two HTH motifs (corresponding to α9 in the predicted structure of BT4124-RR) while two different contact surfaces within the first HTH motif (corresponding to α6-α8 in BT4124-RR) are observed in CdpR and XylR. DNA-recognition helices in these structures are exposed but held at unfavorable positions for binding the target DNA sites. The predicted DBD contact surfaces in HTCS-RRs are distinct from the three described above with the recognition helix α8 buried at the contact surface. Although the domain arrangements predicted by AlphaFold need to be validated by future structural studies, it is unsurprising that the AraC-type DBDs in HTCSs can adopt a different inhibitory contact surface adapted specifically for the REC domain. Given the diversity of interdomain interfaces observed within other RR transcription factor subfamilies (48), the more surprising prediction is that a single domain arrangement appears to be common to most HTCS RRs.

Our mutational studies and structural predictions of HTCS-RRs suggest the α3-β4 loop of the REC domain as the major DBD contact region, providing yet another example for the interaction plasticity of RECs. The α3-β4 loop has been shown to engage in REC-DBD contacts in a few RRs that have DBDs different from the AraC-type DBDs of HTCSs, such as RitR (49), VraR (12) and NarL (50). Analogous to the helix α8 in predicted structures of HTCS-RRs, the DNA recognition helix of NarL is also held at the interdomain surface making extensive contacts with the α2-β3 and α3-β4 loops. Phosphorylation is believed to “open” up the conformation and release the recognition helix for DNA-binding (51, 52).

### Mechanism of phosphorylation-dependent regulation of DBDs

Interdomain contacts of many RRs often involve structural elements in the α4-β5-α5 region of REC. This region has been shown to undergo the largest structural reorganization upon phosphorylation (53–55), as well as to participate in dimerization of RECs, e.g., the α1-α5 dimer in VraR and RcsB (12, 56), and the α4-β5-α5 dimer in the OmpR/PhoB subfamily of RRs (11, 13, 20, 21, 33). REC-DBD interactions not only restrict DBD’s access to DNA but also block dimerization in these RRs to regulate DNA-binding activities. Because covarying residues have been identified in HTCSs at multiple positions corresponding to the α4-β5-α5 dimer interface (14), it is likely RECs in HTCSs can still dimerize. However, it is not clear whether dimerization of RECs within the existing HTCS dimer plays any role in regulation. Unlike the OmpR/PhoB subfamily of RRs in which the α4-β5-α5 region is involved in both REC-DBD interaction and REC dimerization, predicted structures of HTCS-RRs do not show any significant REC-DBD contact at the α4-β5-α5 region, thus the interdomain interaction at the α3-β4 loop is unlikely having a direct impact on dimerization of RECs. Further, other proteins or other HTCS domains that are not investigated in this study may also interact with REC or DBD and contribute to the versatile regulatory strategy commonly seen in TCSs.

The contact residues (S/CH and DK) in HTCSs are suggested to form hydrogen bonds to anchor the REC-DBD interface, which is a common theme observed for RRs. The central question for the regulatory mechanism is how phosphorylation affects these hydrogen bonds to release the DBD. An allosteric mechanism has been described in many RRs involving the phosphorylation-induced rearrangement of T/S at the phosphorylation site correlated with switching of the rotameric orientation of a Y residue in the β5 strand (11, 54, 55). For the OmpR/PhoB subfamily of RRs, such as MtrA (20), PrrA (33) and DrrB (21), the hydrogen bond network at the REC-DBD interface often includes the switch residue Y in β5 and various D or N residues from the DBD. Switching of the Y residue from an outward position to an inward position is believed to disrupt the hydrogen bond and reorient the REC and DBD. This Y residue in β5 is highly conserved in HTCSs (Fig. 4). Even if it is not predicted to interact with the DBD directly, reorientation of the Y sidechain could contribute to conformational changes that allosterically impact the α3-β4 loop. Interestingly, a D residue two positions N-terminal to the Y at the start of β5 is also conserved in HTCSs (Fig. 4). In RR structures that contain a D residue at the same position, the negatively charged side chain forms a salt bridge with a positively charged K or R residue at the end of α3, which happens to be highly conserved in HTCSs as well (Fig. 4). In the predicted HTCS-RR structures, the distance between the conserved D and K residues is within the range of a salt bridge. It is conceivable that phosphorylation-induced structural changes in α4-β5 could propagate to the α3-β4 loop involving this conserved D-K pair. Similarly, deletion of the DBD or amino acid substitutions of the contact residues may also cause changes in the α3-β4 loop propagating through the same allosteric pathway to the phosphorylation active site.

### Conserved and non-consensus contact motifs

The S/CHIP-DK residues at the predicted REC-DBD interface are exceptionally conserved among HTCSs. Percentages of sequence conservation at these positions are 84% (S/C), 91% (H), 96% (I), 98% (P), 95% (D) and 96% (K). This is in stark contrast to residues involved in interdomain interactions between the REC and the histidine kinase HisKA domain, where considerable variations exist. It has been argued that tethering the HK and RR allows a relaxed selection for specific HK-RR interactions (14). The resulting promiscuous phosphotransfer may facilitate rapid expansion of HTCSs for different polysaccharides while a strictly conserved REC-DBD interaction ensures inhibition of the DNA-binding activity during gene duplication and domain shuffling events.

On the other hand, there is still a small fraction of HTCSs with sequences deviating from the conserved S/CHIP-DK. About 6% of HTCSs have an A instead of S/C and 3% have a D or G instead of H at the second position of the motif. In *B. theta*, these HTCSs with deviations from the consensus are predicted to have various domain orientations with little contact, likely due to a loss or decrease in interdomain interaction. This can potentially allow the DBD to bind DNA in the absence of phosphorylation and enable the HTCS to repress gene transcription via binding to a DNA repression site. Indeed, one of these HTCSs, BT2391(<u>F</u>HIP-<u>Q</u>K), has been shown to repress transcription of its corresponding PUL genes (57). Substitutions at the conserved motif in BT4124-RR^AD^ resulted in phosphorylation-independent binding to DNA. Similarly, deviation from the consensus in these HTCSs could also accommodate a regulatory strategy not requiring phosphorylation. BT4236 (C<u>S</u>IP-EK), contains a Q^848^ residue instead of an H at the conserved HK phosphorylation site, suggesting a phosphorylation-independent mechanism. Most of these HTCSs with a non-consensus motif are associated with PULs utilizing host glycans (36, 57).

Host glycans, such as mucin *O*-linked glycans, are less preferred than dietary glycans by *B. theta* and PULs for mucin glycans are usually repressed by other high priority polysaccharides (41, 58). Deviation of the contact residues from the consensus may reflect unique regulatory roles of these HTCSs in host glycan utilization.

Variations in regulatory mechanisms utilized by RR components of HTCSs are unsurprising given the great diversity of regulatory strategies that have been observed for other RRs. While diversity is the norm, common domain arrangements have been noted for some RR subfamilies (11). Our studies have identified a predominant REC-DBD domain arrangement in HTCSs that appears to underlie a fundamental regulatory mechanism. The presence of S/CHIP- DK contact residues is predictive of an interdomain interface that inhibits DNA-binding in the unphosphorylated RR. Disruption of the inhibitory interaction by substitutions of the conserved contact residues can potentially lead to universal activation of individual HTCSs, providing a strategy to facilitate characterization of the direct regulatory targets of HTCSs, especially those whose signals have not been identified.

## Materials and Methods

### Strains and plasmids

Strains and plasmids used in this study are listed in Table S2. *E. coli* strain DH5α was used for cloning. A cloning vector pT7GG2, containing a T7 promoter, a *lac* operator, *lacI^q^*, two *Bsa*I sites and a C-terminal His-tag (GSGAGGHHHHHHG), was constructed by golden-gate assembly of PCR fragments. Different HTCS genes were amplified by PCR using the genomic DNA of *Bacteroides thetaiotaomicron* VPI 5482 (ATCC, 29148D-5) as templates. PCR products were purified and cloned between the two *Bsa*I sites of pT7GG2 by golden-gate cloning. To create corresponding HTCS genes encoding variants with interface substitutions, site-directed mutagenesis was done with specific primers containing the desired mutations and the PCR products were cloned into pT7GG2 similarly as above. All plasmids were confirmed by sequencing.

Promoter regions of *bt4114* and *bt4662* were selected for DNA-binding assays. Both promoters drive transcription of *susCD* homologs of corresponding PULs, thus are potential regulatory targets of BT4124 and BT4663. Binding sites for BT4124 and BT4663 have been predicted within the promoters of *bt4114* and *bt4662*, respectively (9, 59). PCR products of both promoters were further cloned into a vector in which two universal primers, HTCSp-up (5’- FAM-GATGGTAGTGTGGGGTCTCATG) and HTCSp-dn (5’-CCTCCTTATTGAATTTCGGTCTCG) can be used to generate fluorescent DNA fragments for EMSA. Sequences of the DNA fragments used for EMSA are listed in the Supplementary Text.

### Protein purification

Amino acid sequences of purified HTCS proteins are listed in the Supplementary Text. HTCS proteins were expressed from BL21(DE3) containing corresponding plasmids. Overnight cultures were inoculated into LB broth supplemented with 100 μg/ml ampicillin and incubated at 37 °C with shaking. Isopropyl β-d-thiogalactopyranoside was added to a final concentration of 0.5 mM after the optical density reached 0.6. After 3 h induction at 37 °C, cells were harvested by centrifugation and lysed by sonication in sonication buffer, 20 mM Tris pH8.0, 100 mM NaCl, and 5 mM β-mercaptoethanol (BME). Lysates were centrifuged (∼75000×g), filtered and loaded onto a 5-ml HisTrap FF column (Cytiva). Bound his-tagged proteins were eluted with a gradient (20-500 mM) of imidazole-containing buffer. Fractions containing the desired proteins were pooled, concentrated using Amicon Ultra 15-ml centrifugal filters (MilliporeSigma), passed through a 0.2 μm filter, and loaded onto a Superdex 75 26/60 column (GE Healthcare) equilibrated with 20 mM Tris, pH 8.0, 100 mM NaCl, and 5 mM BME. For BT4124-RR proteins, an additional step of purification using an anion exchange HiTrap Q column (Cytiva) equilibrated with 20 mM Tris, pH 8.0, and 5 mM BME and elution with a 0-1 M NaCl gradient, was applied before loading onto the Superdex 75 column. Fractions containing HTCS proteins eluted from the Superdex 75 column were pooled and stored at -80 °C after rapid freezing in a dry ice/ethanol bath.

### Autophosphorylation of HTCS proteins

All phosphorylation reactions were performed with 5 μM HTCS proteins in Tris buffer (50 mM Tris, pH 7.5, 100 mM NaCl, 5 mM BME and 5 mM MgSO_4_). After addition of phosphoramidate (PAM) at specified concentrations, 9-μl aliquots were removed from the reaction mixture at indicated times and immediately mixed with 4×SDS sample loading buffer to stop the reactions. Time intervals were usually 0, 10 s, 20 s, 40 s, 2 min, 10 min and 30 min for proteins with fast phosphorylation kinetics, and 0, 30 s, 60 s, 2 min, 4 min, 10 min and 30 min for proteins with slow kinetics. Aliquots were stored on ice before analysis using phos-tag gels. Phos-tag gels were prepared as described previously with 20-25 μM Phos-tag acrylamide (Wako Chemicals), and 50 μM MnCl_2_ (28, 60). For HTCS-RR proteins, 11% acrylamide was used;13.5% acrylamide was used for HTCS-REC proteins with lower molecular weights. Phos-tag gels were electrophoresed at 170 V at room temperature until dyes entered the wells, then were transferred to ice baths, and the electrophoresis continued at 130 V for 35 min. Proteins were visualized using Coomassie Blue and quantified with ImageJ. The fraction of phosphorylated protein was calculated by measuring the intensity of the upper shifted band (phosphorylated) relative to the total intensity of both protein bands within each lane.

It has been shown that some small-molecule phospho-donors will increase the ionic strength, and can potentially decrease phosphorylation rates, impacting the kinetic analyses (61, 62).

Effects of ionic strength on phosphorylation were evaluated by comparing reactions without ionic strength correction and with salts added to keep a constant ionic strength (see details in Supplementary Text). Our results suggest that the ionic strength corresponding to 50 mM PAM did not significantly impact phosphorylation rates for HTCS proteins (Fig. S1). Phosphorylation kinetics presented in all other figures are from data without correction for ionic strengths.

Kinetics of RR autophosphorylation have been characterized for several RRs by tracking the quenching of the intrinsic tryptophan fluorescence upon phosphorylation and fitting the time course to determine kinetic constants (61–64). The four HTCS-RRs do not contain tryptophan and Phos-tag analyses are limited by the number of time points for a robust fitting. For phosphorylation data (20 mM PAM) that contain considerable time points, phosphorylation fractions from multiple independent experiments were pooled and globally fitted with an exponential function to illustrate the trendline of phosphorylation equilibrium. At other PAM concentrations, phosphorylation data were obtained focusing on the initial stage of phosphorylation (3-4 time points) to calculate the initial rates from a linear regression. To derive the *k_cat_*/*K_m_*values for individual HTCS proteins, initial rates of phosphorylation were fitted with the Michaelis-Menten kinetics. For simplicity, phosphorylation cooperativity of the potential REC or RR dimer is not considered (see details in Supplementary Text). To accommodate the initial rates with units of fraction phosphorylated instead of concentration (conc.), the classic Michaelis-Menten rate equation is rewritten as below,

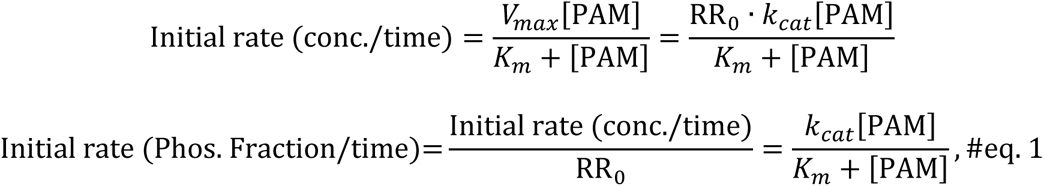

in which *RR_0_* is the total protein concentration. Initial rates at different PAM concentrations from multiple independent experiments were fitted globally with eq. 1. Phosphorylation efficiency was calculated as *k_cat_*/*K_m_*and standard deviation of the efficiency was derived from standard deviations of fitted *k_cat_* and *K_m_* using the standard error propagation formula.

### Structure prediction

Sequences of HTCS proteins from *B. theta* were retrieved from Uniprot(65) and used for structure prediction using AlphaFold2 via the ColabFold platform (30, 31). Sequences from 5-10 residues before the start of the REC domain to the C-terminus of protein were used for HTCS-RRs while sequences from ∼10 residues after the transmembrane region to the C-terminus were used for HTCS-cyto. Monomeric HTCS-RR and dimeric HTCS-cyto structures were predicted with default parameters. Focusing on the REC-DBD domain arrangements, all predicted structures were visualized and analyzed using Pymol. RMSD values were calculated by aligning the predicted RR structures with BT4124-RR. Refinement cycle was set at zero to ensure the entire structure was used for alignment without eliminating highly variable atoms.

### Analyses of sequence conservation in HTCS-RRs

HTCS protein sequences were retrieved from Uniprot by searching proteins containing “histidine kinase”, “response regulatory” and “HTH araC/xylS-type” domains within *Bacteroides* (taxonomy id, 816). The resulting 6908 protein sequences were aligned using MAFFT (66), and the alignment was input to the HMMER package in UGENE (67) to generate the profile hidden Markov model (HMM). HMMs for the REC domain (PFAM id, PF00072) and the DNA-binding HTH18 domain (PFAM id, PF12833) were downloaded from InterPro (68). HMMs were analyzed by Skylign (69) to generate the sequence logo and derive above-background information contents (ICs) at each position. ICs of all amino acids were summed to yield the total IC at each position. Total ICs were normalized by subtracting the background and dividing by the peak IC for comparison of sequence conservation.

### DNA-binding of HTCS-RRs

DNA fragments labeled with 5’-fluorescein were used for EMSAs to assess the DNA-binding activities of HTCS-RRs. For binding assays with phosphorylated proteins, HTCS-RRs were phosphorylated using 50 mM PAM for 45 min prior to the DNA-binding reactions. Proteins of different concentrations were mixed with ∼2.5 ng/μl of DNA in binding buffer (Tris 50 mM, pH 7.6, 200 mM NaCl, 2.5% glycerol [v/v], 2 mM MgCl_2_, and 0.5 mM DTT) containing 0.1 μg/μl bovine serum albumin and 10 ng/μl of salmon sperm DNA. After 30 minutes of incubation, 1.5 μl of loading dye (15% Ficoll 400 [w/v], 0.25% bromophenol blue[w/v]) was added to 10 μl of reaction mixture followed by loading to 5% TBE gels. Gels were electrophoresed on ice at ∼130 v for ∼40 min and visualized by fluorescence imaging using a FluorChem Q (Alpha Innotech).

DNA band intensities were quantified using ImageJ and the band intensity of free DNA with no protein added was considered as the 100% standard. The percentage decreases of free unbound DNA due to protein addition were calculated to represent the percentage of bound DNA. Because proteins were not fully phosphorylated under experimental conditions and the exact phosphorylation levels could be affected by DNA-binding, rigorous analyses considering different contributions from phosphorylated and unphosphorylated proteins were not performed. To compare binding curves for different reactions, binding data from different experiments were pooled and globally fitted with the Hill equation using experimental protein concentrations to derive the apparent DNA affinity. A universal protein concentration series (0, 0.6, 1.2, 1.8, 2.4, 4 and 8 μM) was used for better comparison of DNA-binding activities even though it may not be optimal for fitting extremely weak or extremely strong binding curves. For strong binding with few data points at intermediate binding fractions, the binding cooperativity was set at 2 to derive the binding trendline. For weak binding not reaching the steady state, the maximal binding percentage was arbitrarily set at 100 for the Hill fit to derive the binding trendline.

## Data Availability

Sequences, multiple sequence alignments, and structural models used for analyses are available in Zenodo (https://zenodo.org/records/10962917).

## Acknowledgment

This work was supported by a grant from the National Institutes of Health (R35GM131727).

## Supplementary Figures

FIG S1. Autophosphorylation of HTCS proteins is not significantly affected by ionic strength. Phosphorylation kinetics of BT4124 (A-D), BT4663 (E), BT3334 (F) and BT1754 (G) with no salt addition (-) or with added NaCl to reach a constant ionic strength (+). One representative example of Phos-tag gels at the indicated PAM concentration is shown for each protein. Quantification of Phos-tag gels yielded the phosphorylation time course (B), and the early stage of the time course (B inset) was used to calculate the initial rates (C, D, mid and right panels in E, F and G). Data are shown as mean ± SD from three independent experiments for BT4124 and at least two independent experiments for other HTCSs.

FIG S2. Autophosphorylation kinetics of HTCS-RR and HTCS-REC proteins. (A-C) Phos-tag gel analyses of protein phosphorylation at 20 mM PAM for BT4663, BT3334 and BT1754. One representative example is shown for each protein. The lower panel of (A) shows an example of phosphorylation time course of BT4663 based on Phos-tag gel quantification. (D) Phosphorylation of BT4663 at different PAM concentrations. One representative example is shown for each condition. Longer times were used for BT4663-RR to allow sufficient phosphorylation to be observed and quantified for calculating the initial rates. (E-G) Dependence of phosphorylation initial rates on PAM concentrations. Data are shown as mean ± SD from at least three independent experiments. Lines with corresponding shaded ranges indicate the fitted curves and the 95% confidence intervals. The shaded ranges for BT4633-RR and BT1754-RR are barely noticeable due to a narrow confidence interval.

FIG S3. Structures of HTCS-RRs predicted by AlphaFold 2. (A) Superposition of the five top-ranked BT4124-RR structures. From rank 1 to rank 5, structures are colored violet, light cyan, green, yellow and wheat. All five structures superimpose well with each other, except for helix α4 and the connecting loop (α5-α6) between the REC and DBD domains. (B) BT4124 structure colored with the pLDDT confidence score. The majority of the structure, except for the α5-α6 connecting loop, the N- and C-termini, have high confidence scores above 50. (C) Superposition of the predicted BT4124-RR structure with the DNA-bound HTH18 family member, MarA (1BL0). The domain arrangement in the predicted BT4124-RR structure appears to not allow DNA-binding due to the steric clash of the REC domain with DNA.

FIG S4. Sequence conservation of HTCS-DBD domains. Sequence logos were obtained from the profile hidden Markov model of 6908 HTCS proteins in Bacteroides. Secondary structural elements are shown above the logos. DBD domains of HTCSs belong to the HTH18 family (PFAM id, PF12833). Bar graphs compare the normalized information contents (ICs) between HTCS-DBDs (gray) and the entire HTH18 family (pink). HMM of HTH18 only contains residues starting from helix α7 (vertical dashed line), thus ICs were not calculated for HTH18 preceding α7. Residues involved in the predicted REC-DBD interfaces (red box) appear highly conserved in HTCS-DBDs, but not in the entire HTH18 family.

FIG S5. Predicted structures of HTCSs from B. theta. RMSD values are derived from structural alignment with BT4124-RR. (A) Similar REC-DBD domain orientation for HTCSs containing the conserved S/CHIP-DK motif. Small RMSD values indicate high structural similarity to BT4124. For BT1754, the conserved REC-DBD domain orientation is only observed in the predicted dimeric structure of BT1754-cyto that contains both the HK and RR domains. (B) Diverse domain orientations in HTCSs that have motif sequences that deviate from the consensus. The helix α5 in the REC domain, the DNA recognition helices α8 and α11 are labeled to highlight different orientations of the DBD. All structures shown are the top ranked predictions from AlphaFold. For HTCSs without the consensus motif, low ranked predictions (rank 2-5) also have large RMSD values, suggesting REC-DBD orientations distinct from the conserved one observed in BT4124-RR.

FIG S6. Plasticity of domain orientations for different ranked structural predictions. One representative, BT2628-RR, is shown here with five different DBD orientations for five ranked predictions. All ranked predictions have high pLDDT scores, suggesting well-modelled local structural elements. The pTM scores are at intermediate values of ∼0.5 not dramatically different for five ranked predictions, suggesting the inter-domain accuracy is not high, likely caused by the relative mobility of individual domains.

FIG S7. Autophosphorylation kinetics of BT4124 domains and corresponding interface variants. Initial rates of phosphorylation at different PAM concentrations were measured from phos-tag gels shown in Figure 5. Lines with corresponding shaded ranges indicate the curves fitted with the Michaelis-Menten equation and the 95% confidence intervals. (A) Differences in phosphorylation kinetics are apparent for BT4124-RR and corresponding interface variants, BT4124-RRAG and BT4124-RRAD, suggesting differences in the inhibitory interactions between the REC and DBD domains. (B) BT4124-REC and BT4124-RECAD have similar rate curves with overlapping confidence ranges, suggesting that the substitution in the isolated receiver domain did not alter the phosphorylation efficiency.

FIG S8. Autophosphorylation kinetics of BT4663-RR and corresponding interface variants. Phos-tag gels (A) were quantified to track the fraction of phosphorylated proteins (B) at indicated times after addition of 20 mM PAM. Lines in (B) represent the global exponential fit to illustrate the kinetic trendline of phosphorylation. Initial rates of phosphorylation were measured from early stages of the reaction to derive the kinetic curves in (C). Similar to that observed for BT4124, the interface variants, BT4663-RRAG and BT4663-RRAD, showed faster phosphorylation kinetics than BT4663-RR, suggesting relief of inhibitory interactions.

